# Spontaneous alpha-band lateralization extends persistence of visual information in iconic memory by modulating cortical excitability

**DOI:** 10.1101/2024.10.23.619788

**Authors:** Paul J.C. Smith, Niko A. Busch

## Abstract

Pre-stimulus alpha oscillations in the visual cortex modulate neuronal excitability, influencing sensory processing and decision-making. While this relationship has been demonstrated mostly in detection tasks with low-visibility stimuli, interpretations of such effects can be ambiguous due to biases, making it difficult to clearly distinguish between perception-related and decision-related effects. In this study, we investigated how spontaneous fluctuations in pre-stimulus alpha power affect iconic memory, a high-capacity, ultra-short visual memory store. Data from 49 healthy adults (34 female and 15 male) was analyzed. We employed a partial report task, where a brief display of six stimuli was followed by a report cue indicating the target stimulus. In this paradigm, accuracy at short stimulus-cue onset asynchronies (SOAs) is typically high, reflecting the initial availability of sensory information, but it rapidly declines at intermediate SOAs due to the decay of the iconic memory trace, stabilizing at a low asymptote at long SOAs, representing the limited capacity of short-term memory. Crucially, performance in this task is constrained by the temporal persistence of sensory information, not by low visibility or response bias. We found that strong pre-stimulus alpha power enhanced performance by amplifying initial stimulus availability without affecting the speed of iconic decay. This effect was driven predominantly by stronger pre-stimulus alpha power in the hemisphere ipsilateral to the to-be-reported target, likely suppressing neuronal excitability of neurons coding irrelevant stimuli. Our findings underscore the role of alpha oscillations in modulating neuronal excitability and visual perception, independent of decision-making strategies implicated in prior studies.

## Introduction

One of the most prominent neuronal signals in the human occipital cortex is the alpha rhythm, oscillating at around 8-13 Hz. Spontaneous pre-stimulus alpha power and phase fluctuations are associated with periodic fluctuations in neuronal excitability (Buzsáki & Draguhn, 2004; Haegens et al., 2011; Dougherty et al., 2017). These fluctuations affect visual perception, attention, and metacognition (Ergenoglu et al., 2004; van Dijk et al., 2008; Limbach & Corballis, 2017; Samaha et al., 2017). Furthermore, these effects are often lateralized across the left and right cortical hemispheres. For instance, shifts of spatial attention due to external cues or internal decisions to attend are associated with alpha power increases in the hemisphere ipsilateral to the attended location, indicating relative inhibition of the hemisphere representing the unattended location, while alpha power decreases contralaterally, reflecting greater excitability in the hemisphere representing the attended location (Worden et al., 2000; Thut et al., 2006; Bengson et al., 2014). However, exactly how alpha power and excitability at the time of stimulus onset affect perceptual decision-making is still debated.

Studies testing the effect of pre-stimulus alpha power have demonstrated that weaker power is associated with higher hit rates in the detection of near-threshold stimuli (Ergenoglu et al., 2004; van Dijk et al., 2008). Initially, this was interpreted as an improvement in the accuracy of visual detection; however, subsequent studies showed that the increase in hit rates accompanies an increase in false alarm rates, confidence, and subjective visibility, indicating a more liberal detection criterion (Benwell et al., 2017; Iemi et al., 2017; Samaha et al., 2017; Balestrieri & Busch, 2022). Interpreting these findings within the classical signal detection framework is challenging. A more liberal criterion can reflect a change in the observer’s deliberate decision-making strategy, independent of their subjective perception, or an amplified representation of both signal and noise, giving both a more signal-like subjective appearance (Samaha et al., 2020). Distinguishing between these interpretations is particularly difficult in the context of detection tasks with near-threshold stimuli, in which performance is limited by the discriminability of signal and noise, and observers apply a criterion for reporting the presence or absence of a stimulus. Here, we propose to overcome these problems by focusing on the effect of pre-stimulus alpha power on temporal stimulus availability rather than detectability.

Here, we investigated the previously unexplored effect of alpha oscillations on temporal persistence of visual information in iconic memory—a high-capacity, short-duration visual memory store that follows visual perception and precedes visual short-term memory (Sperling, 1960; Dick, 1974). We tested iconic memory performance and confidence using a partial report paradigm (Figure 1A), in which a display comprising six concentric stimuli was briefly flashed, and the to-be-reported object was indicated by a cue that appeared after a variable stimulus-cue onset asynchrony (SOA; Lu et al., 2000). Performance in this task is excellent when the cue and display are presented simultaneously, implying that all the objects are available for report at cue onset. At longer display-cue SOAs, performance drops only gradually, implying that some stimulus information persists for a few hundred milliseconds after stimulus offset, reflecting the decay of information within a high-capacity iconic memory store (Gegenfurtner & Sperling, 1993). Teeuwen et al. (2021) demonstrated that iconic memory is based on the persistent firing of neurons in the primary visual cortex. Given that the strength of neuronal responses in the early visual cortex is correlated with the alpha rhythm (Dougherty et al., 2017; Lundqvist et al., 2020), we hypothesized that weak pre-stimulus alpha power (reflecting high neuronal excitability) would amplify the neural response or slow down its decay (Figure 1B), thereby extending the persistence of information in iconic memory. Importantly, performance in this paradigm is not limited by sensory noise and is not influenced by a detection criterion. Thus, any effects of pre-stimulus alpha power on temporal persistence would reflect a genuine effect on perception, rather than strategic decision-making.

**Figure 1:**
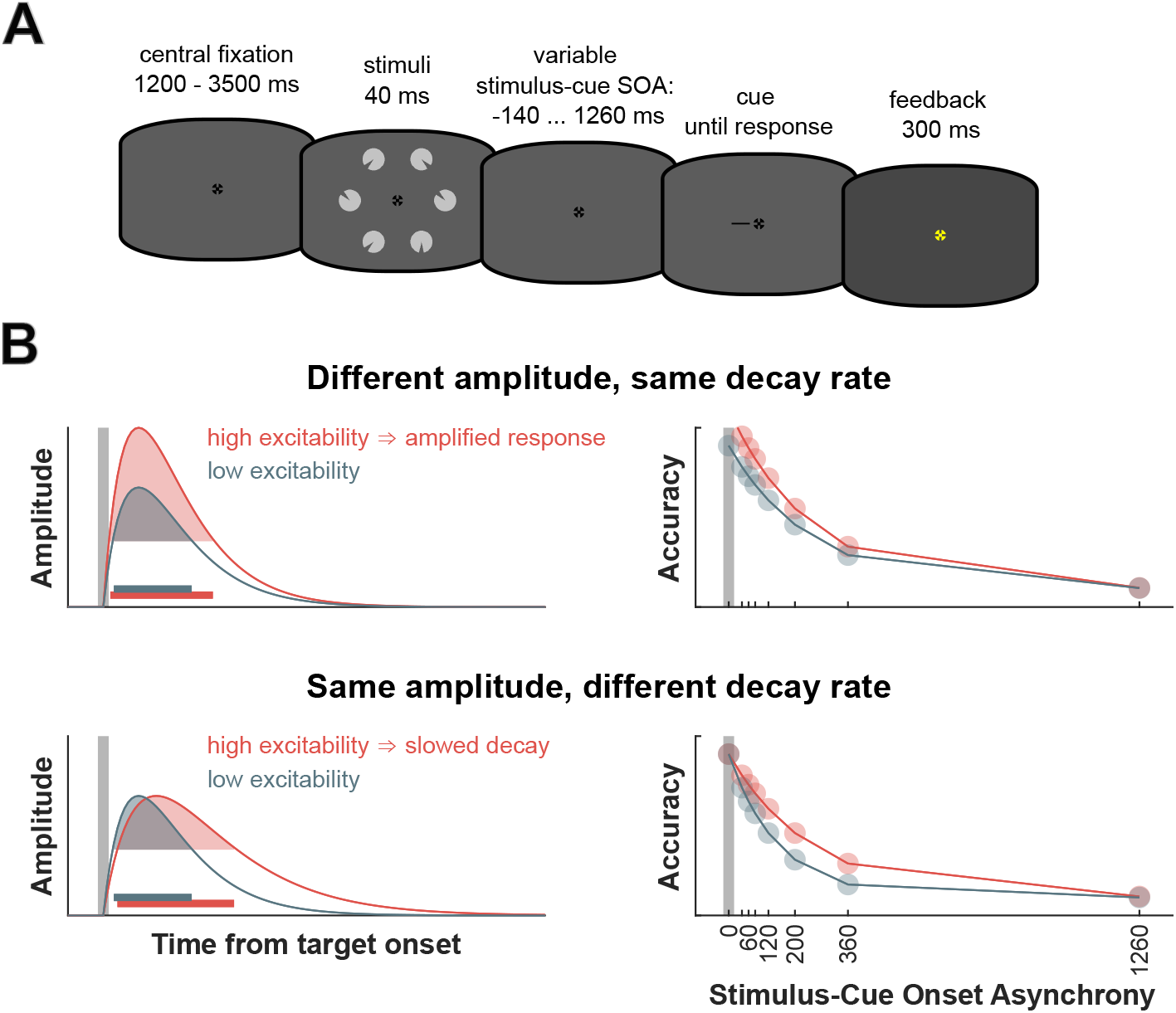
**A**: Illustration of the partial report paradigm. Each trial began with a central fixation cross displayed for a variable interval of 1200-3500 ms. Following fixation, a stimulus array consisting of six items was presented for 40 ms. Participants were then cued to report the orientation of one target item, with the cue appearing at various SOAs relative to stimulus onset: 140 ms before stimulus onset, at stimulus onset, or 40, 60, 80, 120, 200, 360, or 1240 ms after. There was no time limit for reporting or the subsequent confidence rating. After the confidence rating, feedback was given via the color of the fixation cross, turning blue for a correct response and yellow for an incorrect one. **B**: Illustration of two possible mechanisms of temporal persistence modulation. Left: Schematic representation of the impulse response triggered by stimulus onset (indicated by the vertical gray bar). The response shows a rapid onset followed by a gradual decay. The stimulus information persists as long as the response amplitude remains above a critical threshold (shaded area). Horizontal bars indicate duration of persistence. Heightened excitability can extend this persistence either by increasing the *amplitude* of the initial response (top) or by slowing its *decay* (bottom). Right: Hypothetical performance in the partial report task as a function of stimulus-cue SOA. Performance is high at short SOAs and declines with longer SOAs. An amplified response (top) is predicted to enhance initial stimulus availability, thereby boosting performance primarily at short SOAs, which would be captured by the model’s *a*_1_ parameter (see equation 1). A slower decay (bottom) would improve performance at intermediate SOAs, reflected in the model’s *τ* parameter.

## Material and Methods

### Pre-registration

The study was pre-registered with the Open Science Framework (osf.io/6rwvt/), and we adhered to the outlined methods unless stated otherwise.

### Participants

We collected EEG and eye-tracking data from 61 healthy participants (aged 23.59 ± 3.94 years), of whom 45 were female and 16 were male. We collected more participants than stated in the preregistration due to favorable conditions. All participants had normal or corrected-to-normal vision, no reported history of neurological or psychiatric disorders, provided their written consent, and were compensated with either course credits or money. The ethics commission of the University of Münster approved the study (ref. 2023-27-PS). One participant was excluded due to poor data quality, five additional participants due to subpar performance (less than 80% accuracy at -140 and 0 ms stimulus onset asynchrony (SOA)), and a further six participants because their performance data could not be fit with the exponential decay model (see below). The final sample size for the main analysis was 49 (aged 23.35 ± 4.1 years), with 34 female and 15 male participants, corresponding to the desired sample size stated in the pre-registration. For the lateralization-based analyses, 12 more participants were excluded due to not completing enough trials, leaving the sample size for this analysis at 37 (aged 23.32 ± 4.38 years): 23 female and 14 male.

### Stimuli and Procedure

The experiment was presented on a 24-inch Viewpixx/EEG LCD Monitor with a 120 Hz refresh rate, 1 ms pixel response time, 95% luminance uniformity, and 1920×1080 pixels resolution (33.76×19.38; www.vpixx.com). The recording took place in a dimly lit, soundproof cabin. Participants’ heads were stabilized on a chinrest with their eyes approximately 86 cm from the monitor. Eye movements were monitored using a desktop-mounted Eyelink 1000+ infrared-based eye-tracking system (SR Research Ltd.) set to a 1000 Hz sampling rate (monocular, from the participant’s dominant eye).

The stimuli used in the partial-report paradigm were circles with wedges cut out at variable orientations (0°, 45°, 90°, 135°, 180°, 225°, 270°, or 315°). The circles had a diameter of 2.6 degrees visual angle (dva) whilst the cut-out had a thickness of 0.28 dva with a diameter of 0.6 dva. The circles were light grey (RGB: [180 180 180]), and the cut-out wedges were of a darker grey (RGB: [70 70 70]), matching the background color. The cue utilized was a black line (RGB: [0 0 0]) with a length of 1 dva and a thickness of 0.1 dva, pointing to one of the former positions of the stimuli. Participants were instructed to avoid eye movements and blinks during the stimulus presentation. Participants had to fixate on the center of the screen, where a black (RGB: [0 0 0]) fixation cross with a size of 0.6 dva was presented for a variable time interval of 1200 to 3500 ms. Following this interval, six stimuli were arranged in a circle around the fixation cross and presented for 40 ms. The cue appeared after a variable SOA of either 140 ms before stimuli onset, at stimuli onset, or 40, 60, 80, 120, 200, 360, or 1240 ms after stimuli onset. Participants were instructed to press a number on the number pad corresponding to the orientation of the wedge in the target circle (eight for 0°, six for 90°, two for 180°, etc.). Following this decision, the participants were instructed to press either four (low), five (medium), or six (high) on the number pad, indicating their respective confidence rating. There was no time limit for their response. Feedback on their decision was provided 300 ms after their confidence rating, with correct responses marked by a blue (RGB: [0 0 255]) fixation cross, and incorrect responses indicated by a yellow (RGB: [255 255 0]) fixation cross. A visualization of a trial can be seen in Figure 1. There were 1296 trials, with short self-paced breaks after 170 consecutive trials to counterbalance SOA, stimulus position around the fixation cross, and target cut-out orientation. The experiment was written and presented using Matlab2022 (mathworks.com), and Psychtoolbox (Brainard, 1997; Pelli, 1997; Kleiner et al., 2007).

### Modeling the Time Course of Iconic Memory

This study aimed to test the hypothesis that pre-stimulus alpha power and neuronal excitability modulate the temporal availability of stimulus information. We assumed that the response to a brief stimulus follows an impulse response, which can outlast the stimulus itself due to the low-pass filtering properties of the early visual system (Dilollo 1980, Loftus & Irwin, 1998).The longer the response amplitude remains above a task-specific critical threshold (shaded areas in Figure 1B), the longer the stimulus information persists, remaining available for perceptual or decision-making processes. Greater neuronal excitability may enhance persistence in two ways: by amplifying the response amplitude and initial sensory availability (Figure 1B, top), or by slowing the response’s decay (Figure 1B, bottom). A similar concept has been proposed to explain the effects of oscillatory frequency on the temporal integration of double flashes (Karvat & Landau 2024). To directly test the effect of alpha power on stimulus persistence and distinguish between its impact on response amplitude versus decay speed, we employed a partial report task. A display of six items was followed by a cue indicating the target to be reported (Figure 1A). Intuitively, performance at short target-cue SOAs reflects the initial availability and response amplitude, while the performance decline across intermediate SOAs reveals the speed of iconic decay.

To quantify these effects and parameters formally, we analyzed the behavioral data using a signal detection theory framework, calculating *d*^*′*^ for each SOA. Behavioral data (*d*^*′*^ and confidence) was fitted to a non-linear decay function,

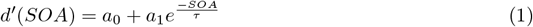

using a nonlinear least-squares solver with start parameters based on Lu et al. (Lu et al., 2005). This analysis was independent of the EEG data. The function consisted of three free parameters: *a*_0_, which refers to the sensitivity at long delays that reflects the amount of information transferred into short-term memory without the cueing benefit; *a*_1_, which refers to the fast-decaying sensitivity that represents the initial visual availability of stimulus information; and *τ*, which refers to the time-constant of this decay. This function was fit to performance data from all non-negative stimulus-cue SOAs. Participants with significant deviations in these parameters were excluded from the analysis (see Participants). Confidence data from all non-negative stimulus-cue SOAs was normalized to a 0-1 scale and then fitted to the same decay function using free-floating start parameters. Participants with significant deviations in the function parameters were excluded from the subsequent modeling.

### EEG acquisition and pre-processing

EEG activity was recorded using a Biosemi Active Two EEG system with 67 Ag/AgCl electrodes (BioSemi B.V.) set to a 1024 Hz sampling rate. Sixty-four electrodes were arranged in a custom-made montage with equidistant placement (EASYCAP GmbH) with additional external electrodes placed next to the right eye, the left eye, and below the left eye.

All EEG data preprocessing and analysis steps were scripted, and run in Matlab2023a (mathworks.com) using the EEGLAB toolbox (Delorme & Makeig, 2004), and custom scripts. The continuous data was downsampled to 256 Hz and re-referenced to the average. It was then high-pass filtered at 0.1 Hz (preregistered was 0.5 Hz), and low-pass filtered at 40 Hz. Following this, the data was epoched from -1000 ms to 1500 ms time-locked to stimulus onset (preregistered was -400 ms to 200 ms). A round of trial rejection based on thresholding (± 500 µV), and joint probability (function pop_jointprob in EEGLAB, local threshold 9, global threshold 5) was performed. Furthermore, trials in which eyeblinks were detected in a range of -500 ms to 500 ms around stimulus onset, or in which participants significantly deviated from the fixation cross (2.5 dva) were rejected. After this, ICA was applied, and components were identified by the IClabel algorithm (Pion-Tonachini et al., 2019) as “Brain”, “Muscle”, “Eye”, “Heart”, “Line Noise”, “Channel Noise”, or “Other”. Components were manually screened, and components classified as non-brain activity were excluded. Following this, noisy channels (SD *>* 2) were spherically interpolated.

### EEG Data Analysis

Pre-stimulus spectral EEG power was computed using a Fast Fourier Transform (FFT) of the data within the pre-stimulus time range from -500 ms to -2 ms before stimulus onset for all electrodes and frequencies. In deviation from the pre-registration, we did not perform an analysis using the FOOOF algorithm (Donoghue et al., 2020) due to time constraints. Our analysis was focused on power averaged across electrodes corresponding approximately to P5, P3, P4, and P6 in the frequency range of 8-12 Hz. Single-trial power was then sorted into three bins using quantile ranking. Performance and confidence data were averaged across the trials in each bin for each non-negative stimulus-cue SOA. The exponential decay model was then fit to d’ and confidence data separately for each bin. At this step, the start parameters of the decay function were chosen based on the best-fitting parameters of the global function fit described above.

In an exploratory analysis, we investigated the effect of pre-stimulus lateralization on performance. The rationale was that a spontaneous relative increase in alpha power in one hemisphere might impede the processing of stimuli represented in that hemisphere, i.e. of stimuli appearing in the contralateral hemifield. Such inhibition would reduce performance on trials where the upcoming cue required reporting a stimulus from the contralateral hemifield, and aid performance when an ipsilateral stimulus was cued. To this end, spectral power was quantified separately for trials with correct and incorrect responses to cued stimuli in the left and right hemifield based on the FFT analysis described above and on a time-frequency analysis using Morlet wavelets. A lateralization index was computed by subtracting contralateral pre-stimulus alpha power relative to the to-be cued stimuli from ipsilateral pre-stimulus alpha power relative to the to-be cued stimuli and normalizing this by dividing it through the sum of both (Thut et al., 2006): *latindex* 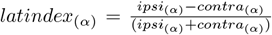 . Lateralized pre-stimulus alpha power was determined based on average power at custom electrode locations corresponding approximately to P3 (left) and P4 (right) in the frequency range of 8-12 Hz. This lateralization index was then binned using quantile ranking into three bins. *d*^*′*^ was computed for each bin and each non-negative stimulus-cue SOA, and the data was again fitted to the decay function. Based on this binning, confidence data was similarly fitted to the decay function using the global confidence decay function parameters. The same binning and decay function fitting was used separately for ipsi-, and contralateral alpha relative to the to-be-cued stimuli. This was done on the performance and the confidence data, both using the best-fitting parameters of their respective global function fit.

### Statistical Analysis

The decay function parameters *a*_0_, *a*_1_, and *τ*, were compared using t-tests between the weakest and strongest non-lateralized pre-stimulus alpha bins, as were the parameters of the weakest and strongest lateralization index bins, ipsilateral bins, and contralateral bins. In deviation from the pre-registration, we decided against using a jackknife approach as left-out single trials make vastly different contributions to the different model parameters depending on the trials’ SOA.

As alpha lateralization has been associated with eye movements and miniature gaze shifts (van Ede et al., 2019; Popov et al., 2023, Mössing et al., 2024), we tested whether pre-stimulus alpha lateralization was confounded by pre-stimulus gaze shifts. Specifically, we tested whether correct responses to targets on the left or right side were preceded by pre-stimulus gaze displacements to the left or right of central fixation. To this end, we used a 2×2 repeated measures analysis of variance (ANOVA) with cue direction (left/right) and correctness (correct/incorrect) as fixed factors, subject id as a random factor, and gaze shift in dva relative to the fixation cross in the interval from 500ms to 2ms before stimulus onset as dependent variable.

### Data and code accessibility

The data will be available upon acceptance of the manuscript at the Open Science Framework (osf.io/xfkrm/). The code will be available at github.com/pauljcs/alpha-iconic.

## Results

### Performance Across All Conditions

#### Bilateral Pre-stimulus Alpha Power

Initial stimulus availability (*a*_1_) was higher for strong pre-stimulus alpha power than weak pre-stimulus alpha power (*t*(48) = −3.82, *p <* 0.001). The function parameters representing the capacity for information transferred to short-term memory (*a*_0_) and the time constant representing the speed of iconic decay (*τ*) did not show significant differences between trials with weak vs. strong pre-stimulus alpha power (*a*_0_ : *t*(48) = 0.53, *p* = 0.6; *τ* : *t*(48) = 0.16, *p* = 0.88; Figure 2A).

**Figure 2:**
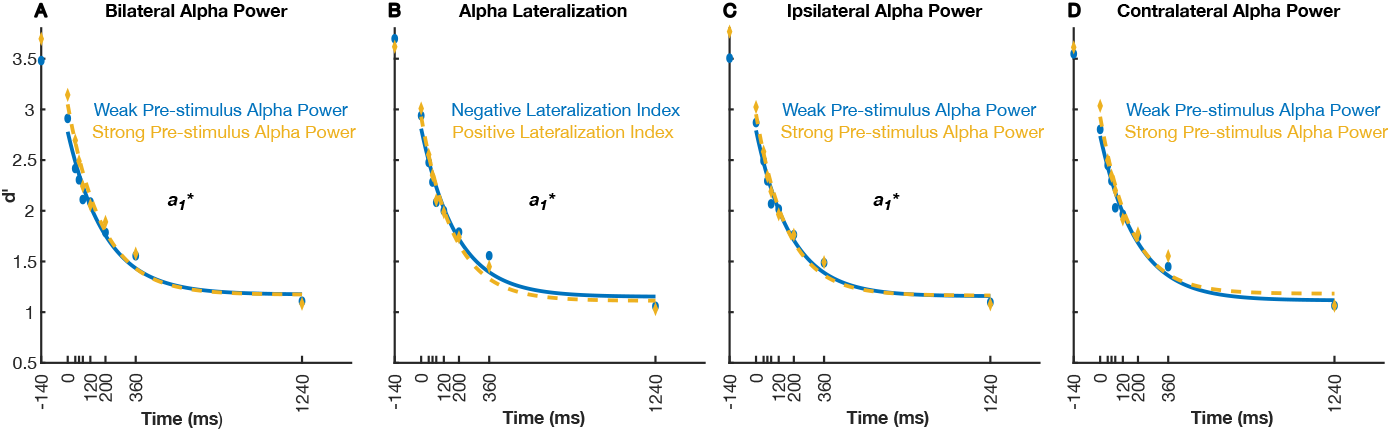
Accuracy (*d*^*′*^) data for each SOA. Lines show model fits. Data are shown separately for: (A) trials with strong vs. weak pre-stimulus alpha power; (B): positive vs. negative pre-stimulus lateralization, where positive lateralization implies lateralization “towards’ the location of the upcoming target item; (C): strong vs. weak pre-stimulus power at ipsilateral channels, relative to the upcoming, cued target item; (D) strong vs. weak pre-stimulus power at contralateral channels. Model parameters showing a significant difference between these conditions are indicated with an asterisk.

For illustration purposes, we complemented this analysis by plotting the difference in power between correct and incorrect trials, irrespective of SOA. This analysis confirmed that trials with correct responses were preceded by stronger pre-stimulus alpha-band power (Figure 4A).

**Figure 3:**
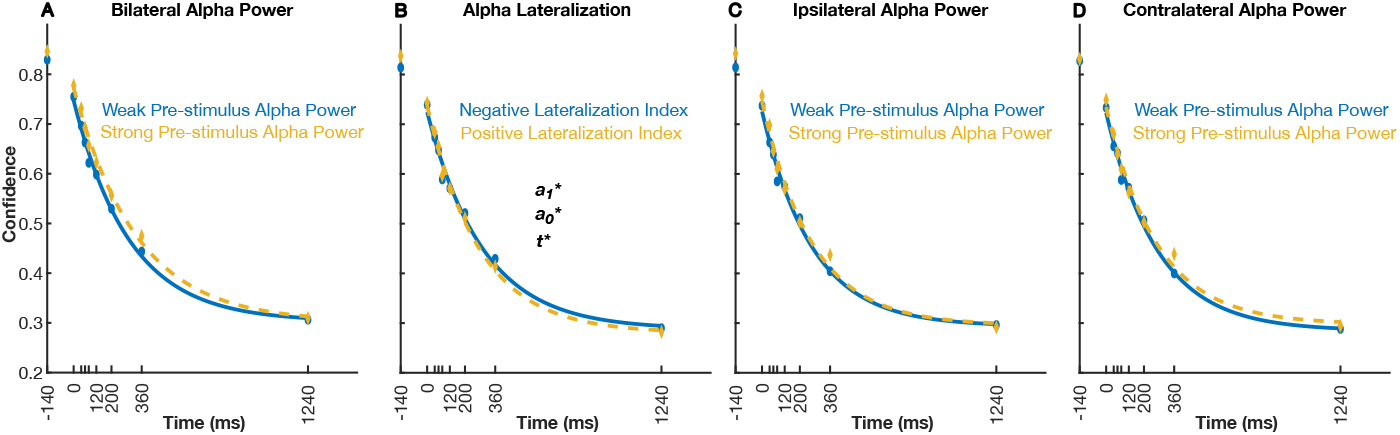
Normalized confidence data for each SOA. Lines show model fits. Data are shown separately for: (A) trials with strong vs. weak pre-stimulus alpha power; (B): positive vs. negative pre-stimulus lateralization, where positive lateralization implies lateralization “towards’ the location of the upcoming target item; (C): strong vs. weak pre-stimulus power at ipsilateral channels, relative to the upcoming, cued target item; (D) strong vs. weak pre-stimulus power at contralateral channels. Model parameters showing a significant difference between these conditions are indicated with an asterisk.

**Figure 4:**
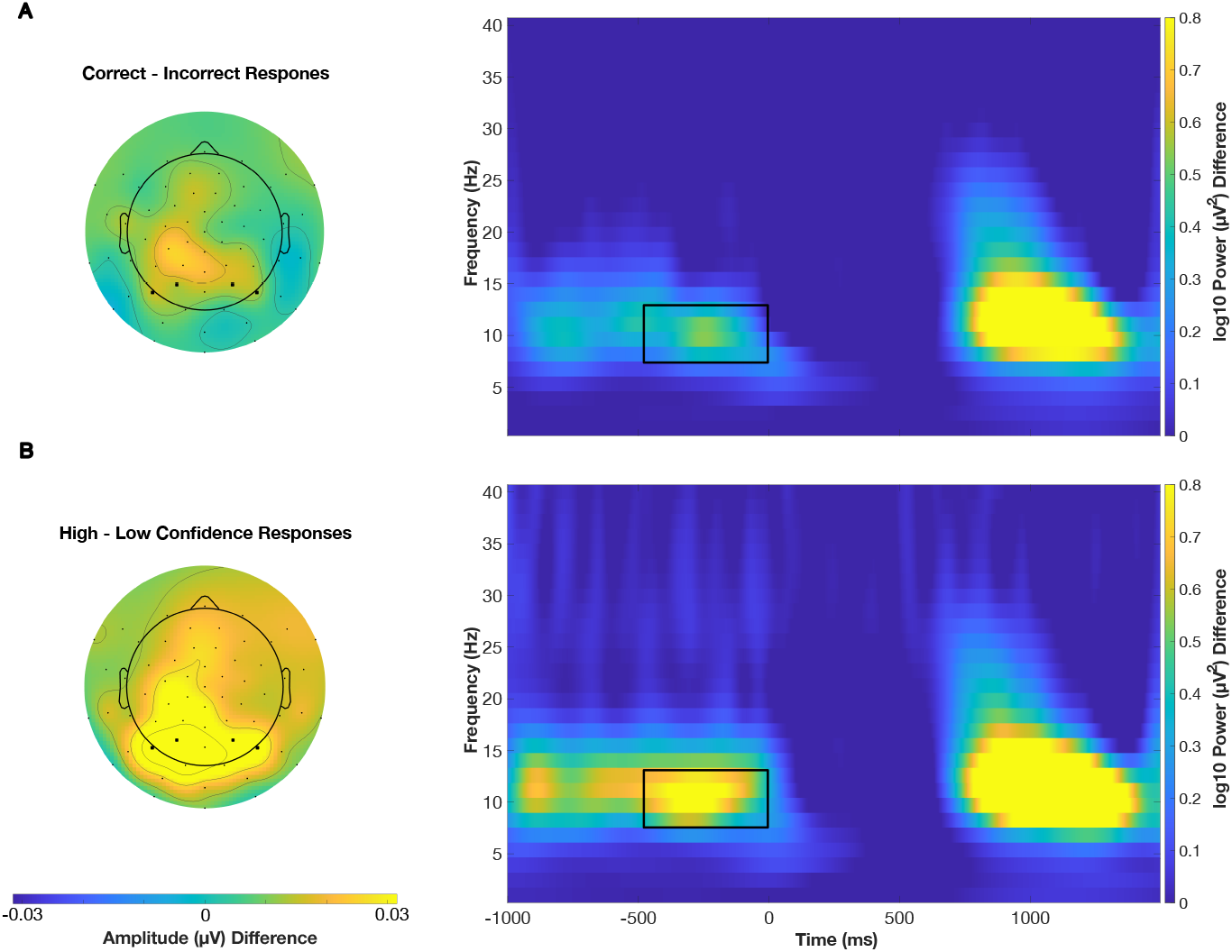
(A) Topography and time frequency plot showing the difference between correct and incorrect responses. Topographies represent the time-frequency window indicated by the black rectangle. Time-frequency plots represent the electrodes highlighted by the bold markers on the topography. (B) Topography and time frequency plot showing the difference between high and low confidence responses. Conventions as in A.

#### Lateralized Pre-stimulus Alpha Power

The parameter representing the initial stimulus availability (*a*_1_) was significantly higher for positive lateralization than negative lateralization (*t*(36) = −2.32, *p* = 0.026). This indicates that performance was better if the ipsilateral to the to-be cued stimuli pre-stimulus power was stronger relative to the contralateral to the to-be cued stimuli pre-stimulus power. However, *a*_0_(the transfer to short-term memory) and *τ* (the speed of iconic decay) did not show significant differences (*a*_0_ : *t*(36) = 0.9, *p* = 0.375; *τ* : *t*(36) = 0.08, *p* = 0.94) for positive and negative lateralization (Figure 2B).

We complemented this analysis by plotting the difference between the correct vs. incorrect effect for target items on the left from that for items on the right. This analysis confirmed a positive difference, reflecting stronger lateralization “towards” the upcoming cued item on correct trials. (Figure 5A).

**Figure 5:**
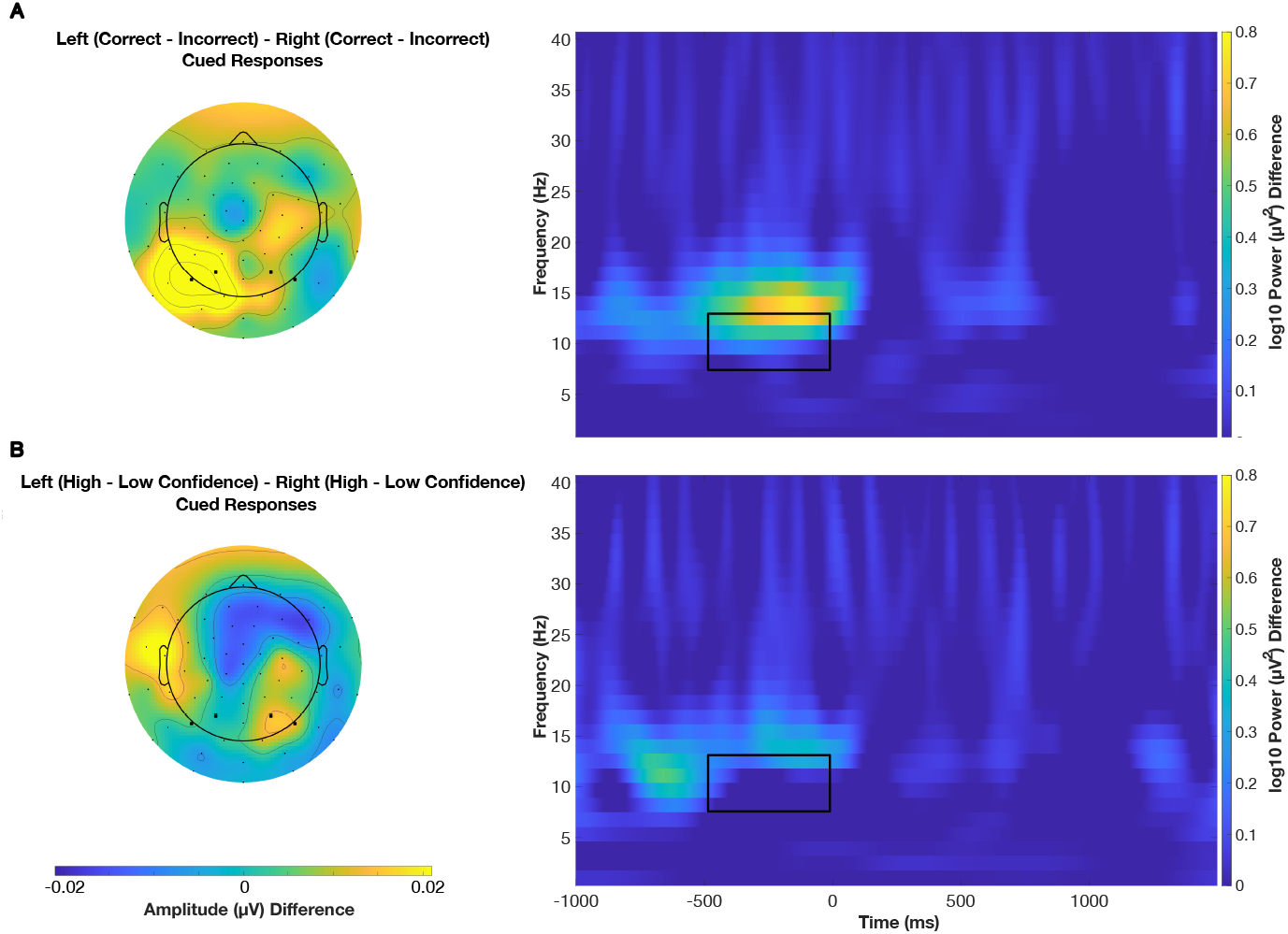
(A) Topography and time frequency plot showing the difference between correct and in-correct responses to target items on the left vs. correct and incorrect responses to right items. Topographies represent the time-frequency window indicated by the black rectangle. Time-frequency plots represent the electrodes highlighted by the bold markers on the topography. (B) Topography and time frequency plot showing the difference between high and low confidence responses to target items on the left vs. high and low confidence responses to right items. Conventions as in A.

To determine whether this effect is driven by a) strong ipsilateral pre-stimulus power or b) weak contralateral pre-stimulus power, we performed the same analysis separately for both ipsi- and contralateral pre-stimulus power.

#### Ipsilateral Pre-stimulus Alpha Power

A significant difference was found for initial stimulus availability (*a*_1_) (*t*(36) = −2.41, *p* = 0.021) between weak and strong ipsilateral pre-stimulus alpha power, with strong ipsilateral pre-stimulus power resulting in better performance. However, no significant differences were found for the parameters representing the transfer to short-term memory (*a*_0_) or the speed of iconic decay (*τ*) (*a*_0_ : *t*(36) = −0.62, *p* = 0.539; *τ* : *t*(36) = 1.11, *p* = 0.275; 2C).

#### Contralateral Pre-stimulus Alpha Power

No significant differences were found for the initial stimulus availability (*a*_1_), transfer to short-term memory (*a*_0_), or speed of iconic decay (*τ*) (*a*_1_ : *t*(36) = −1.53, *p* = 0.134; *a*_0_ : *t*(36) = −1.65, *p* = 0.107; *τ* : *t*(36) = 1.89, *p* = 0.067) between weak and strong contralateral pre-stimulus alpha power (2D). Weaker contralateral pre-stimulus power appears to not affect iconic memory performance whereas stronger ipsilateral pre-stimulus power does, thus indicating that the effect found for spontaneous lateralization does result from stronger ipsilateral pre-stimulus power.

### Confidence Across All Conditions

#### Bilateral Pre-stimulus Alpha Power

The function parameters representing initial stimulus availability (*a*_1_), information transfer to short-term memory (*a*_0_), and speed of iconic decay (*τ*) did not show any significant differences (*a*_1_ : *t*(48) = −0.7, *p* = 0.485; *a*_0_ : *t*(48) = 0.7, *p* = 0.487; *τ* : *t*(48) = −0.79, *p* = 0.434) for weak vs. strong pre-stimulus alpha power (Figure 3A).

For illustration purposes, we complemented this analysis by plotting the difference in power between high and low confidence trials, irrespective of SOA. This analysis confirmed that trials with high confidence responses were preceded by stronger pre-stimulus alpha-band power (Figure 4B).

#### Lateralized Pre-stimulus Alpha Power

Significant differences were found for initial stimulus availability (*a*_1_), transfer to short-term memory (*a*_0_), and speed of iconic decay (*τ*) (*a*_1_ : *t*(36) = 13.37, *p <* 0.001; *a*_0_ : *t*(36) = 12.84, *p <* 0.001; *τ* : *t*(36) = −2.93, *p* = 0.006) for negative and positive lateralization (Figure 3B). Positive lateralization results in better performance, memory transfer and slower decay of iconic memory. Again, we performed the same analysis separately for ipsi- and contralateral pre-stimulus alpha power in order to determine the cause of this effect.

We complemented this analysis by plotting the difference between the high vs. low confidence effect for target items on the left from that for items on the right. This analysis confirmed a weak, but positive difference, reflecting stronger lateralization “towards” the upcoming cued item on high confidence trials. (Figure 5B).

#### Ipsilateral Pre-stimulus Alpha Power

No significant differences were observed for the function parameters representing initial stimulus availability (*a*_1_), information transfer to short-term memory (*a*_0_), and speed of iconic decay (*τ*) (*a*_1_ : *t*(36) = −1, *p* = 0.324; *a*_0_ : *t*(36) = 1, *p* = 0.325; *τ* : *t*(36) = −1, *p* = 0.325) for weak vs. strong ipsilateral pre-stimulus alpha power (Figure 3C).

#### Contralateral Pre-stimulus Alpha Power

No significant differences were found for the function parameters representing initial stimulus availability (*a*_1_), information transfer to short-term memory (*a*_0_), and speed of iconic decay (*τ*) (*a*_1_ : *t*(36) = 0.88, *p* = 0.387; *a*_0_ : *t*(36) = −1.05, *p* = 0.299; *τt*(36) = 0.56, *p* = 0.576) between weak and strong contralateral pre-stimulus alpha power (Figure 3D).

### Gazeshift

The previous analyses showed that participants responded more accurately and with higher confidence when pre-stimulus alpha power was spontaneously lateralized “towards” the position of the upcoming item that would be cued for report. Given that alpha lateralization has been associated with miniature eye movements and gaze drifts (van Ede et al., 2019; Popov et al., 2023, Mössing et al., 2024), we tested if small incidental horizontal gaze displacements before stimulus onset were systematically associated with better performance when the gaze shift was in the direction of the upcoming cued target item. Specifically, we tested if gaze position was systematically displaced to the left on trials with a correct response to a left target, and vice versa. This interaction between correctness and cue direction was not significant, implying that accuracy was not confounded by gaze shifts (*F* (1) = 2.94, *p* = 0.0968). The main effect of correctness (*F* (1) = 1.27, *p* = 0.269) and direction (*F* (1) = 0.21, *p* = 0.649) were also not significant.

## Discussion

How does the ongoing state of neuronal excitability influence sensory information processing? In the present study, we addressed this question by investigating the role of pre-stimulus alpha-band lateralization in modulating the temporal persistence of visual information in iconic memory. Specifically, we hypothesized that spontaneous alpha oscillations modulate the persistence of sensory information by altering cortical excitability, thereby modulating iconic memory performance.

This hypothesis was grounded in the theoretical framework suggesting that alpha oscillations reflect ongoing changes in neuronal excitability, which have been shown to influence perceptual decision-making. Numerous studies (reviewed in Samaha et al., 2020) have demonstrated that near-threshold targets are less likely to be detected during periods of high alpha power, reflecting reduced excitability (Ergenoglu et al., 2004; Van Dijk et al., 2008). Importantly, these effects are due to a more conservative detection criterion rather than impaired accuracy (Limbach & Corballis, 2016; Iemi et al., 2017). As a result, pre-stimulus alpha power has a much less pronounced effect in tasks that do not rely on detection criteria, such as feature discrimination tasks (Iemi et al., 2017). However, signal detection theory does not fully explain how changes in decision criteria arise, and the theoretical implications of these findings remain debated (Samaha et al., 2022). One interpretation is that pre-stimulus brain states reflect a decision bias, a post-perceptual adjustment in decision-making independent of subjective perception (Kloosterman et al., 2019). Another interpretation suggests a perceptual bias, where the intensity and subjective signal-likeness of sensory input and noise are modulated (Samaha et al., 2020; Benwell et al., 2022).

To address this ambiguity, we employed a criterion-free iconic memory task, where performance is constrained by the temporal persistence of sensory information rather than near-threshold stimulus intensity. Iconic memory, a high-capacity but short-lived storage system, temporarily holds visual information for a few hundred milliseconds before transferring some content to more durable, but low-capacity short-term memory (Sperling, 1960; Neisser, 1967). It is supported by persistent firing in early visual areas (Duysens et al., 1985; Keysers et al., 2005; Teeuwen et al., 2021), where strong alpha power has been linked to reduced visual-evoked responses (Iemi et al., 2019; Lundqvist et al., 2020). Since the strength and duration of stimulus-evoked responses are critical determinants of iconic memory performance (Di Lollo, 1977, see Figure 1B), we predicted that strong alpha power, reflecting reduced excitability, would shorten stimulus availability and impair partial report performance.

Contrary to our hypothesis, we found that stronger bilateral pre-stimulus alpha power was associated with better partial report performance at short target-cue SOAs, suggesting amplified initial stimulus availability (Figure 2 A; Figure 4A). No effects were observed on model parameters related to the speed of decay or transfer to short-term memory (Figure 2A). Confidence ratings were also unaffected by pre-stimulus alpha power (Figure 3A; Figure 5B), contrasting with previous studies linking stronger alpha power to lower confidence in detection tasks (Benwell et al., 2017; Samaha et al., 2017).

To better understand this unexpected result, we conducted an exploratory analysis of pre-stimulus alpha lateralization, examining power separately in the left and right channels and testing their effects on left and right targets independently. We found that stronger pre-stimulus alpha power specifically improved performance at electrodes ipsilateral to the upcoming target (Figure 2C). For example, stronger alpha power in the left posterior channels enhanced performance for left targets but not for right targets, and a similar pattern was observed for confidence (Figure 2B; Figure 3B; Figure 5B). In contrast, no lateralization effects were found at contralateral channels (Figure 2D). This suggests that the observed association between strong bilateral alpha power and improved performance did not result from power over both hemispheres, but rather from the superposition of two genuinely lateralized effects: strong left-lateralized power enhancing performance for left targets, and right-lateralized power enhancing performance for right targets.

This finding aligns well with research on cue-induced spatial attention, where alpha power typically increases over the hemisphere ipsilateral to a cue and decreases contralaterally. This pattern reflects greater inhibition in the hemisphere processing the irrelevant hemifield and heightened excitability in the hemisphere processing the relevant hemifield (Foxe & Snyder, 2011; Händel et al., 2011). Notably, in our study, pre-stimulus alpha lateralization was not externally driven by an attentional cue but occurred spontaneously, likely due to trial-by-trial fluctuations in self-initiated endogenous attention (Bengson et al., 2014; Nadra et al., 2023) or spontaneous variations in neuronal excitability (Balestrieri & Busch, 2022).

Thus, the analysis of pre-stimulus lateralization supports the hypothesis that strong lateralized pre-stimulus alpha power enhances performance specifically for targets in the ipsilateral hemifield. Moreover, this effect was linked to the model parameter representing initial stimulus availability, reflecting the strength of the stimulus-evoked response, which is essential for iconic memory, visual persistence and temporal integration (Di Lollo, 1977; Karvat & Landau, 2024; Figure 1B, top panel). The lateralized effect may reflect alpha-induced inhibition of the non-cued hemifield, reducing competition during stimulus encoding. Additionally, the absence of effects on the transfer of information to short-term memory suggests that alpha lateralization influences performance by modulating the availability of stimulus information, not by selective encoding into a more stable, capacity limited memory store.

Together, these results demonstrate that spontaneous fluctuations in the momentary state of neurophysiological excitability and inhibition can modulate the temporal availability of sensory information. This supports previous models linking alpha oscillations to perceptual bias (Samaha et al., 2020) and suggests that the detection of near-threshold targets may involve not only amplified stimulus-evoked responses but also prolonged availability due to more persistent responses.

Several avenues for future research arise from these findings. First, our pre-registered analysis of bilateral power produced the unexpected result of strong alpha power being associated with improved performance. While our exploratory follow-up analysis strongly suggests that this reflects the superposition of well-known lateralized effects, which align well with the existing literature on cue-induced and spontaneous lateralization, this interpretation raises new questions. Many previous studies reporting that strong bilateral alpha power impairs detection have used lateralized target stimuli (Chaumon & Busch., 2014; Iemi et al., 2017; Pilipenko & Samaha, 2024). It remains unclear why these effects were not similarly lateralized in those studies. It is conceivable that the lateralization effect was more pronounced in the present study because it used a supra-threshold stimulus display and a task where performance was not limited by low stimulus visibility. However, only a direct comparison between detection and partial report tasks could help resolve this issue. Second, alpha power had a weaker influence on confidence in our study compared to previous findings. This may be due to the high signal-to-noise ratio in our paradigm, where confidence is primarily driven by decision accuracy rather than uncertainty, as in near-threshold paradigms (Peters, 2022). Third, does the lateralized effect of alpha power on iconic memory performance stem from self-initiated spatial attention, and how does it compare to cue-induced attention effects? The relationship between attention and iconic memory has been debated: some authors argue that iconic memory requires little to no attention (Bachmann & Aru, 2016; Aru & Bachmann, 2017; Pinto et al., 2017), while others contend that iconic memory depends on attention (Mack et al., 2015; Mack et al., 2016), or is at least modulated by it (Botta et al., 2019). Therefore, comparing the effects of spontaneous lateralization on iconic memory to those of cue-induced lateralization would offer valuable insights not only into the role of the alpha rhythm but also into the broader theories of iconic memory, attention, and awareness.

## Conclusion

Our study demonstrated that pre-stimulus alpha power influences iconic memory performance through spontaneous lateralization. Strong pre-stimulus alpha power ipsilateral to the to-be-cued stimuli suppresses neuronal excitability in visual areas representing task-irrelevant information, thereby extending the temporal availability of relevant information and improving accuracy. These findings suggest that alpha-induced modulations of excitability not only affect the detection of near-threshold stimuli but also enhance the persistence of visual information in iconic memory. This lateralized effect mirrors the role of cue-induced spatial attention in alpha modulation, contributing to the ongoing discussion on the relationship between attention and iconic memory. Ultimately, our work advances the understanding of how fluctuations in neuronal excitability shape both perception and sensory memory.

## Acknowledgments

We thank Teresa Berther, Milena Carolin Koch, Seray-Ezgi Öztekin, Mathilde Maria Pöppelmann, Johanna Seroka, Simon Steibel, and Charlotte Wellmann for help with the data acquisition.

